# A highly accurate model for screening prostate cancer using propensity index panel of ten genes

**DOI:** 10.1101/2021.03.22.436371

**Authors:** Shipra Jain, Kawal Preet Kaur Malhotra, Sumeet Patiyal, Gajendra P. S. Raghava

## Abstract

Prostate-specific antigen (PSA) is a key biomarker, which is commonly used to screen patients of prostate cancer. There is a significant number of unnecessary biopsies that are performed every year, due to poor accuracy of PSA based biomarker. In this study, we identified alternate biomarkers based on gene expression that can be used to screen prostate cancer with high accuracy. All models were trained and test on gene expression profile of 500 prostate cancer and 51 normal samples. Numerous feature selection techniques have been used to identify potential biomarkers. These biomarkers have been used to develop various models using different machine learning techniques for predicting samples of prostate cancer. Our logistic regression-based model achieved highest AUROC 0.91 with accuracy 82.42% on validation dataset. We introduced a new approach called propensity index, where expression of gene is converted into propensity. Our propensity-based approach improved the performance of classification models significantly and achieved AUROC 0.99 with accuracy 96.36% on validation dataset. We also identified and ranked selected genes which can be used to discriminate prostate cancer patients from health individuals with high accuracy. It was observed that single gene-based biomarkers can only achieve accuracy around 90%. In this study, we got best performance using a panel of 10 genes; random forest model using propensity index.

**Highlights:**

- Application of Machine learning techniques to identify Biomarkers for PRAD cancer.
- Highly accurate models developed for classifying prostate cancer vs. normal sample.
- Introducing Propensity index concept for enhancing model performance.
- Top 10 genes identified using feature selection techniques.

## Introduction

Prostate Adenocarcinoma (PRAD) is the second most prevalent cancer diagnosed in men around the world [1]. Patients with prostate cancer are diagnosed at an advanced stage, as patients hardly develop any symptoms at an early stage. Better understanding of the molecular insights responsible for the onset of prostate carcinogenesis, would help in exploring novel therapeutics methods. In the literature, Prostate specific antigen (PSA) test is a widely used test for detecting prostate cancer at a clinically significant stage for better treatment outcomes [2]. Higher PSA levels could indicate benign prostatic enlargement at an early stage. Due to false positive prediction by this test, it leads to many unnecessarily biopsies. Thus, there is a need to identify novel prostate cancer specific biomarkers [3]. Recently two urine based RNA biomarkers prostate cancer antigen 3 (PCA3) [4] and fusion of two genes TMPRSS2:ERG [5] have also been reported which can be used to distinguish between men with early stage disease from men in higher risk stage. Studies have reported the molecular insights involved in development of prostate adenocarcinoma such as members of the E26 transformation-specific (ETS) family of transcription factors fusions with androgen-regulated promoters (e.g. TMPRSS2) [6] and occurrence of point mutations of TP53, FOXA1, PTEN and SPOP gene [7]. PCA3 (originally named as DD3) is a urine based biomarker, which is widely used for prostate cancer detection [8]. Apart from genomic changes, epigenetic level changes have also been reported in cases of prostate cancer such as GSTP1 hypermethylation reported in up to 70 percent of cases [9].

In one study, researchers claimed to identify a three gene panel (HOXC6, TDRD1, and DLX1) as a promising tool to distinguish men with prostate cancer even though they have been reported with low sPSA values [10]. Researchers proposed a method SelectMDx which analyses RNA based biomarkers HOXC6 and DLX1 via reverse transcription, to reduce the need of initial biopsy test [11]. This method is applied on post-DRE patients, measures the HOXC6 & DLX1 mRNA levels [12]. In one study, researchers have proposed ConfirmMDx method which is a tissue-based epigenetic test, developed in a study of 350 men with negative biopsy or repeat biopsy in last two years [13]. The test builds on a “field effect” phenomenon [14]. Due to limited data and samples available, this method is not regularly recommended in clinical practice.

In the recent studies over better cancer clinical management, use of machine learning techniques have contributed in early detection of cancer disease [15]. There is need of identify reliable biomarkers to for screening of prostate cancer in order to avoid unnecessary biopsies [16]. This motivated us to design this study for identifying biomarkers for screening prostate cancer patients with high precision. In this study, we aimed to identify gene expression-based biomarkers to distinguish between prostate cancer patients and a healthy control. In order to select relevant features, we introduce single-gene based feature selection techniques. These techniques allow to rank genes based on their discrimination power. We select top 10 genes using each feature selection technique. These genes are based on difference in mean, significance difference in mean and area under receiver operating characteristic curve (AUROC). We used seven machine learning techniques to develop prediction model using selected genes for identification of prostate cancer patients. In order to improve the performance, we used propensity index based approach for developing prediction models using propensity instead of expression of genes. This propensity based improve the performance of models significantly.

## Materials and Methods

### Dataset

We downloaded GDC TCGA Prostate Cancer (PRAD) dataset from Xena Browser (https://xenabrowser.net/datapages/) that contain gene expression profile of 500 prostate cancer samples and 51 normal samples. It contains expression of 20530 genes for each sample. In this study, FPKM values of RNA transcripts are used as quantification values. Due to large variation in FPKM value, we normalized values using log2 after addition of 1.0 as a constant number to each of FPKM values.

### Feature selection techniques

In this study we have used three types of feature selection techniques, which are based on Mean, Significance difference in mean and AUROC. We have applied these approaches to identify genes whose expression could easily distinguish between prostate and non-prostate subjects. We have extracted top 10 genes using these feature selection approaches, from a list of 20530 gene identifiers. Following is brief description of each technique.

### Mean based approach

In this approach we have calculated the mean expression of each gene prostate cancer patients as well as for health samples. Then we compute difference between mean expression in prostate cancer and healthy samples for each gene. If difference is high, it means that gene can be used to discriminate two types of samples. We ranked genes based difference in mean; and selected top genes which have maximum difference. Following formula has been used for computing difference in mean for a given gene

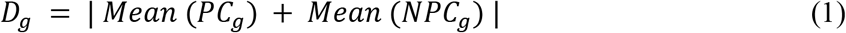

where *D_g_* is difference in mean for gene *g*, *PC_g_* is gene expression of gene *g* for prostate cancer samples, *NPC_g_* is gene expression of gene *g* for non-prostate cancer samples.

Gene identifiers were sorted in decreasing order of the absolute difference in mean values. Top 10 genes identifiers were selected from the sorted list with the highest mean difference between prostate cancer and non-prostate cancer samples.

### Significance difference in mean

In this approach, we compute the level of significance in mean expression of a gene in prostate and non-prostate cancer samples. In addition to mean, we also compute standard deviation in expression of a gene in prostate and non-prostate cancer samples. Following formula is used to compute significance difference in mean for given gene

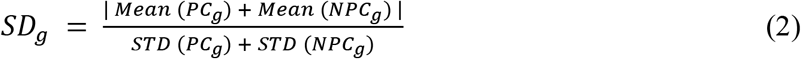

Where *SD_g_* is significance difference in mean of gene *g* in prostate and non-prostate cancer samples. *PC_g_* is gene expression of gene *g* for prostate cancer samples, *NPC_g_* is gene expression of gene *g* for non-prostate cancer samples. *STD* is standard deviation; *STD(PC_g_)* is standard deviation in expression of gene *g* in prostate cancer samples. *STD(NPC_g_)* is standard deviation in expression of gene *g* in non-prostate cancer samples.

The genes are sorted in decreasing order of *SD_g_*, i.e. value calculated by dividing mean by standard deviation. Top 10 genes identifiers were selected from the sorted list with the highest difference between prostate and non-prostate cancer samples.

### Area under curve

In this feature selection technique, we compute the discrimination power of each gene in term AUROC. First of all we calculated the mean of each gene ID for Prostate cancer and non-cancer data respectively. The classification of samples is performed based on expression of a given gene is above or below the threshold value. The threshold value is varied to compute the AUROC from the curve between true positive rate and false positive rate. This process is performed for all genes in dataset. Finally, top 10 genes were selected which have maximum discrimination power in term of AUROC.

**Figure 1:**
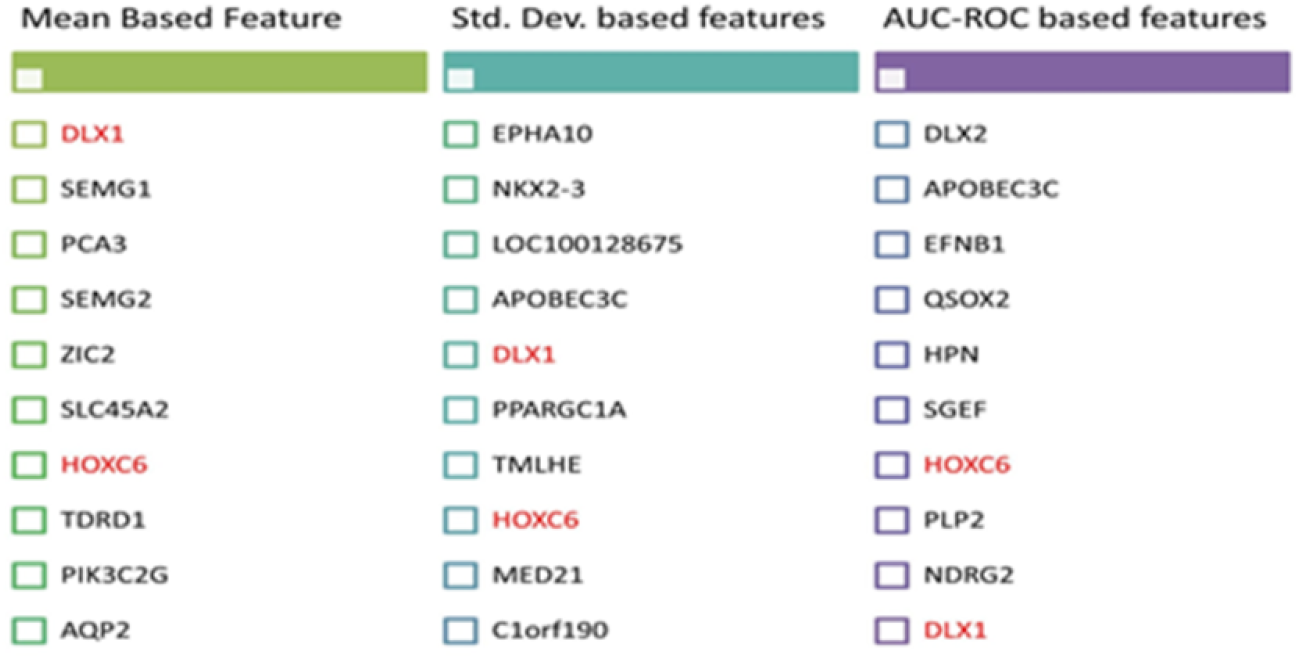
List of genes selected based on different feature selection techniques; top 10 genes from each technique.

### Propensity Index Matrix

In this study, we have coined the concept of propensity index matrix, for feature extraction from gene expression. In this method, we have computed the range of gene expression for a given gene in our dataset and difference has been divided into 10 equal bins. In each bin we compute propensity of prostate and non-prostate cancer samples in each bin. In next step, we replaced the expression value of a gene by a propensity score based on bin it belongs. A new data set is created using propensity index score, which is provided as an input file to machine learning techniques for classification models.

### Application on Machine learning techniques

In this study, we have applied seven different machine learning techniques for developing classification models. These techniques are Support Vector Machine, K-Nearest Neighbour, Decision Tree, Random Forest, Linear Regression, Gaussian Naive Bayes, XGBoost machine learning to our dataset. Using these techniques, we have developed our classification models. These techniques have been implemented using a python library scikit-learn.

### Evaluation of models

In this study, we have used cross validation techniques to evaluate the performance of our models. We divided our dataset randomly into two datasets in the ratio of 70:30, where 70% of data is used for training and 30% of dataset is used for validation. We trained and tested our models on training dataset using five-fold cross-validation technique; where four folds are used as training dataset and remaining one-fold as testing data set. This process of dividing training and testing dataset is repeated five times. The performance evaluation of developed models on the testing dataset is called internal validation. In order to optimize the performance of our models on training dataset we optimized parameters. Final optimized model, best performance in internal validation was used to test on independent or validation dataset.

In order to measure the performance of our models, we used standard parameters commonly used to measure the performance of classification models. Both threshold-dependent and threshold-independent parameters are reported to evaluate the performance. We computed sensitivity, specificity, accuracy and Matthew’s correlation coefficient (MCC) as threshold-dependent parameters using the following equations:

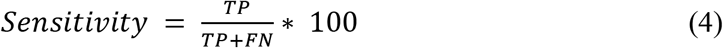

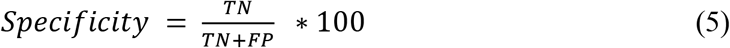

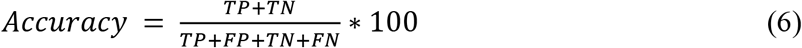

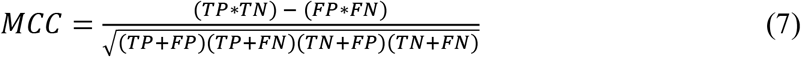

Where, FP is false positive, FN is false negative, TP is true positive and TN is true negative, respectively.

Area Under the Receiver Operating Characteristic curve (AUROC) is reported as a standard parameter for threshold-independent measures.

## Results

We developed classification models for classifying prostate and non-prostate cancer samples using seven machine learning techniques. First, we identified to 10 genes (DLX1, SEMG1, PCA3, SEMG2, ZIC2, SLC45A2, HOXC6, TDRD1, PIK3C2G, and AQP2) using mean-based feature selection techniques. These selected genes were used to build machine learning techniques-based classification models. The performance of all models is evaluated on training and validation dataset. As shown in Table 1, our logistic regression-based model achieved maximum AUROC 0.91 on training as well as on validation dataset.

**Table 1:** The performance of machine learning techniques based models developed using top 10 genes selected using mean based approach.

| Model | Training Dataset |  |  |  |  | Validation Dataset |  |  |  |  |
| --- | --- | --- | --- | --- | --- | --- | --- | --- | --- | --- |
|  | Sens | Spec | Accuracy | AUROC | MCC | Sens | Spec | Accuracy | AUROC | MCC |
| **GNB** | 97.41 | 83.33 | 96.10 | 0.90 | 0.78 | 94.63 | 68.75 | 92.12 | 0.82 | 0.59 |
| **KNN** | 98.85 | 75.00 | 96.63 | 0.87 | 0.79 | 97.32 | 75.00 | 95.15 | 0.86 | 0.73 |
| **SVM** | 94.82 | 88.88 | 94.29 | 0.92 | 0.73 | 85.91 | 75.00 | 84.85 | 0.80 | 0.45 |
| **DT** | 98.56 | 83.33 | 96.36 | 0.90 | 0.78 | 97.99 | 62.50 | 94.54 | 0.80 | 0.66 |
| **RF** | 99.42 | 72.22 | 96.62 | 0.86 | 0.80 | 97.98 | 62.50 | 94.54 | 0.80 | 0.66 |
| **XGB** | 98.56 | 72.22 | 96.10 | 0.85 | 0.76 | 95.95 | 62.50 | 92.73 | 0.79 | 0.58 |
| **LR** | 93.96 | 88.89 | 93.50 | 0.91 | 0.70 | 83.22 | 75.00 | 82.42 | 0.91 | 0.70 |

Similarly, we extract top 10 genes (EPHA10, NKX2-3, LOC100128675, APOBEC3C, DLX1, PPARGC1A, TMLHE, HOXC6, MED21, C1orf190) using significance difference in mean-based feature selection techniques. These to 10 genes were used to build classification models using machine learning techniques. The performance of these models were evaluated on training and validation/testing dataset. As shown in Table 2, our Support vector machine (SVM) based model achieved highest AUROC 0.92 on training dataset and AUROC 0.89 on validation datasets.

**Table 2:** The performance of machine learning techniques based models developed using top 10 genes selected using significance difference in mean.

| Model | Training Dataset |  |  |  |  | Validation Dataset |  |  |  |  |
| --- | --- | --- | --- | --- | --- | --- | --- | --- | --- | --- |
|  | Sens | Spec | Accuracy | AUROC | MCC | Sens | Spec | Accuracy | AUROC | MCC |
| **GNB** | 97.41 | 91.97 | 96.88 | 0.95 | 0.83 | 95.30 | 81.25 | 93.94 | 0.88 | 0.70 |
| **KNN** | 98.56 | 86.11 | 97.40 | 0.92 | 0.85 | 95.30 | 62.50 | 92.12 | 0.79 | 0.56 |
| **SVM** | 95.40 | 88.89 | 94.80 | 0.92 | 0.74 | 90.60 | 87.50 | 90.30 | 0.89 | 0.62 |
| **DT** | 97.70 | 77.77 | 96.09 | 0.88 | 0.75 | 97.32 | 56.25 | 93.33 | 0.76 | 0.58 |
| **RF** | 97.98 | 77.78 | 96.35 | 0.88 | 0.77 | 97.31 | 75.00 | 95.15 | 0.86 | 0.72 |
| **XGB** | 97.99 | 80.56 | 96.36 | 0.89 | 0.79 | 96.64 | 81.25 | 95.15 | 0.89 | 0.74 |
| **LR** | 95.40 | 97.22 | 95.57 | 0.96 | 0.80 | 91.95 | 87.50 | 91.15 | 0.88 | 0.65 |

Finally, we used AUROC based approach for feature selection, where top ten genes were selected based on their performance. There genes are (DLX2, APOBEC3C, EFNB1, QSOX2, HPN, SGEF, HOXC6, PLP2, NDRG2, DLX1) shown highest performance in term of AUROC when we used threshold-based model for prediction. As shown in Table 3, our K-means nearest neighbor (KNN) based model obtained maximum AUROC 0.92 on training dataset and AUROC 0.91 and testing datasets.

**Table 3:** The performance of machine learning techniques based models developed using top 10 genes selected using AUROC based approach.

| **Model** | **Training Dataset** |  |  |  |  | **Validation Dataset** |  |  |  |  |
| --- | --- | --- | --- | --- | --- | --- | --- | --- | --- | --- |
|  | **Sens** | **Spec** | **Accuracy** | **AUROC** | **MCC** | **Sens** | **Spec** | **Accuracy** | **AUROC** | **MCC** |
| **GNB** | 94.54 | 94.45 | 94.53 | 0.94 | 0.75 | 93.29 | 87.50 | 92.73 | 0.90 | 0.68 |
| **KNN** | 98.27 | 86.11 | 97.14 | 0.92 | 0.83 | 95.30 | 87.50 | 94.55 | 0.91 | 0.74 |
| **SVM** | 90.51 | 94.45 | 90.90 | 0.92 | 0.65 | 88.59 | 87.50 | 88.48 | 0.88 | 0.58 |
| **DT** | 97.99 | 83.34 | 96.10 | 0.91 | 0.80 | 96.64 | 68.75 | 93.94 | 0.83 | 0.65 |
| **RF** | 98.85 | 72.22 | 97.40 | 0.86 | 0.77 | 97.31 | 81.25 | 95.76 | 0.89 | 0.76 |
| **XGB** | 98.27 | 75.00 | 96.09 | 0.87 | 0.76 | 97.31 | 81.25 | 95.76 | 0.89 | 0.76 |
| **LR** | 92.53 | 94.45 | 92.72 | 0.93 | 0.70 | 88.59 | 87.50 | 88.48 | 0.88 | 0.58 |

### Models based on propensity Index

In this study, we added a new concept for developing classification models. Instead of using expression of a gene as input, we used propensity of a gene as input. In order to convert expression of a gene to propensity index of a gene, we divide range of expression in 10 bins. In next step, we compute propensity index for each bin. Finally, expression of a gene is converted into a propensity based it expression fall in to a given bin. We developed classification models using top 10 genes selected by mean based feature selection techniques. As shown in Table 4, we got maximum AUROC 1.0 on training dataset and 0.91 on testing dataset using Random Forest (RF) model. The performance of our models improved significantly when we used propensity index instead of expression (See Table 1 and 4).

**Table 4:** The performance of different machine learning techniques based models developed using top 10 genes selected by mean based features selection technique. The models were developed using propensity index of genes instead of their expression.

| Model | Training Dataset |  |  |  |  | Validation Dataset |  |  |  |  |
| --- | --- | --- | --- | --- | --- | --- | --- | --- | --- | --- |
|  | Sens | Spec | Accuracy | AUROC | MCC | Sens | Spec | Accuracy | AUROC | MCC |
| **GNB** | 98.50 | 100.00 | 98.64 | 1.00 | 0.93 | 100.00 | 80.00 | 98.18 | 0.90 | 0.89 |
| **KNN** | 98.50 | 100.00 | 98.64 | 1.00 | 0.93 | 100.00 | 80.00 | 98.18 | 0.89 | 0.89 |
| **SVM** | 98.75 | 97.62 | 98.64 | 1.00 | 0.93 | 100.00 | 60.00 | 96.36 | 0.90 | 0.76 |
| **DT** | 99.25 | 92.86 | 98.64 | 0.96 | 0.92 | 100.00 | 30.00 | 93.64 | 0.65 | 0.53 |
| **RF** | 98.50 | 100.00 | 98.64 | 1.00 | 0.93 | 100.00 | 70.00 | 97.27 | 0.91 | 0.82 |
| **XGB** | 97.75 | 97.62 | 97.74 | 0.99 | 0.88 | 100.00 | 80.00 | 98.18 | 0.93 | 0.89 |
| **LR** | 98.75 | 100.00 | 98.87 | 1.00 | 0.94 | 100.00 | 70.00 | 97.27 | 0.87 | 0.82 |

Similarly, we developed models based on propensity index of 10 gene selected using significance difference in based feature selection. As shown in Table 5, logistic regression based (LR) based model achieved highest performance with AUROC 1.00 on training dataset and AUROC 0.97 on validation dataset. In comparison to table 2 statistics, performance of prediction models reported in table 5 have increased after converting expression values to propensity index values.

**Table 5:** The performance of different machine learning techniques based models developed using top 10 genes selected by significance difference in mean based approach. The models were developed using propensity index of genes instead of their expression.

| Model | Training Dataset |  |  |  |  | Validation Dataset |  |  |  |  |
| --- | --- | --- | --- | --- | --- | --- | --- | --- | --- | --- |
|  | Sens | Spec | Accuracy | AUROC | MCC | Sens | Spec | Accuracy | AUROC | MCC |
| **GNB** | 99.50 | 100.00 | 99.55 | 1.00 | 0.97 | 100.00 | 70.00 | 97.27 | 0.90 | 0.82 |
| **KNN** | 99.75 | 100.00 | 99.77 | 1.00 | 0.99 | 100.00 | 70.00 | 97.27 | 0.95 | 0.82 |
| **SVM** | 99.00 | 100.00 | 99.10 | 1.00 | 0.95 | 100.00 | 80.00 | 98.18 | 0.95 | 0.89 |
| **DT** | 97.75 | 90.48 | 97.06 | 0.94 | 0.84 | 99.00 | 60.00 | 95.45 | 0.80 | 0.69 |
| **RF** | 99.75 | 95.24 | 99.32 | 1.00 | 0.96 | 100.00 | 80.00 | 98.18 | 0.94 | 0.89 |
| **XGB** | 98.75 | 97.62 | 98.64 | 1.00 | 0.93 | 99.00 | 80.00 | 97.27 | 0.93 | 0.83 |
| **LR** | 99.00 | 100.00 | 99.10 | 1.00 | 0.95 | 100.00 | 80.00 | 98.18 | 0.97 | 0.89 |

Finally, we developed models using propensity index of top 10 genes obtained from AUROC based feature selection approach. As shown in Table 6, Random Forest (RF) model obtain best performance with AUROC 1.00 on training dataset and 0.99 on validation dataset. It is clear from above results that performance models developed using propensity index (Table 4, 5, 6) got better performance than models developed using gene expression (Table 1, 2, 3).

**Table 6:** The performance of different machine learning techniques based models developed using top 10 genes selected by AUROC based approach. The models were developed using propensity index of genes instead of their expression.

| Model | Training Dataset |  |  |  |  | Validation Dataset |  |  |  |  |
| --- | --- | --- | --- | --- | --- | --- | --- | --- | --- | --- |
|  | Sens | Spec | Accuracy | AUROC | MCC | Sens | Spec | Accuracy | AUROC | MCC |
| **GNB** | 99.50 | 100.00 | 99.55 | 1.00 | 0.97 | 100.00 | 50.00 | 95.45 | 0.85 | 0.69 |
| **KNN** | 99.50 | 100.00 | 99.55 | 1.00 | 0.97 | 100.00 | 50.00 | 95.45 | 0.84 | 0.69 |
| **SVM** | 99.25 | 100.00 | 99.32 | 1.00 | 0.96 | 100.00 | 70.00 | 97.27 | 0.95 | 0.82 |
| **DT** | 97.75 | 90.48 | 97.06 | 0.94 | 0.84 | 99.00 | 60.00 | 95.45 | 0.80 | 0.69 |
| **RF** | 98.25 | 100.00 | 98.42 | 1.00 | 0.92 | 99.00 | 70.00 | 96.36 | 0.99 | 0.76 |
| **XGB** | 98.50 | 95.24 | 98.19 | 0.99 | 0.90 | 99.00 | 80.00 | 97.27 | 0.97 | 0.83 |
| **LR** | 98.75 | 100.00 | 98.87 | 1.00 | 0.94 | 100.00 | 70.00 | 97.27 | 0.78 | 0.82 |

### Single gene based classification

In order to understand importance of individual gene in discriminating prostate and non-prostate samples. We developed threshold-based models that can be used to identify prostate cancer samples based on expression of a single gene. Thus, after identifying best genes using feature selection techniques, we have also ranked them on the basis of their capability of correctly predicting prostate cancer samples. All 30 gene extracted using feature selection techniques (i.e. top 10 from mean-based method, 10 from standard deviation based and 10 from AUROC based method) are considered for ranking. After removing duplicate genes, we ranked these genes based on probability of correct prediction of prostate cancer samples. As shown in figure 2, we got 13 genes which have probability of correct prediction from 0.97 to 0.989. In figure 2, we also added the performance of KLK3 a gene associated with PSA (commonly use test). It is clear that the performance of KLK3 gene is too poor in comparison to other genes used in our study.

**Figure 2:**
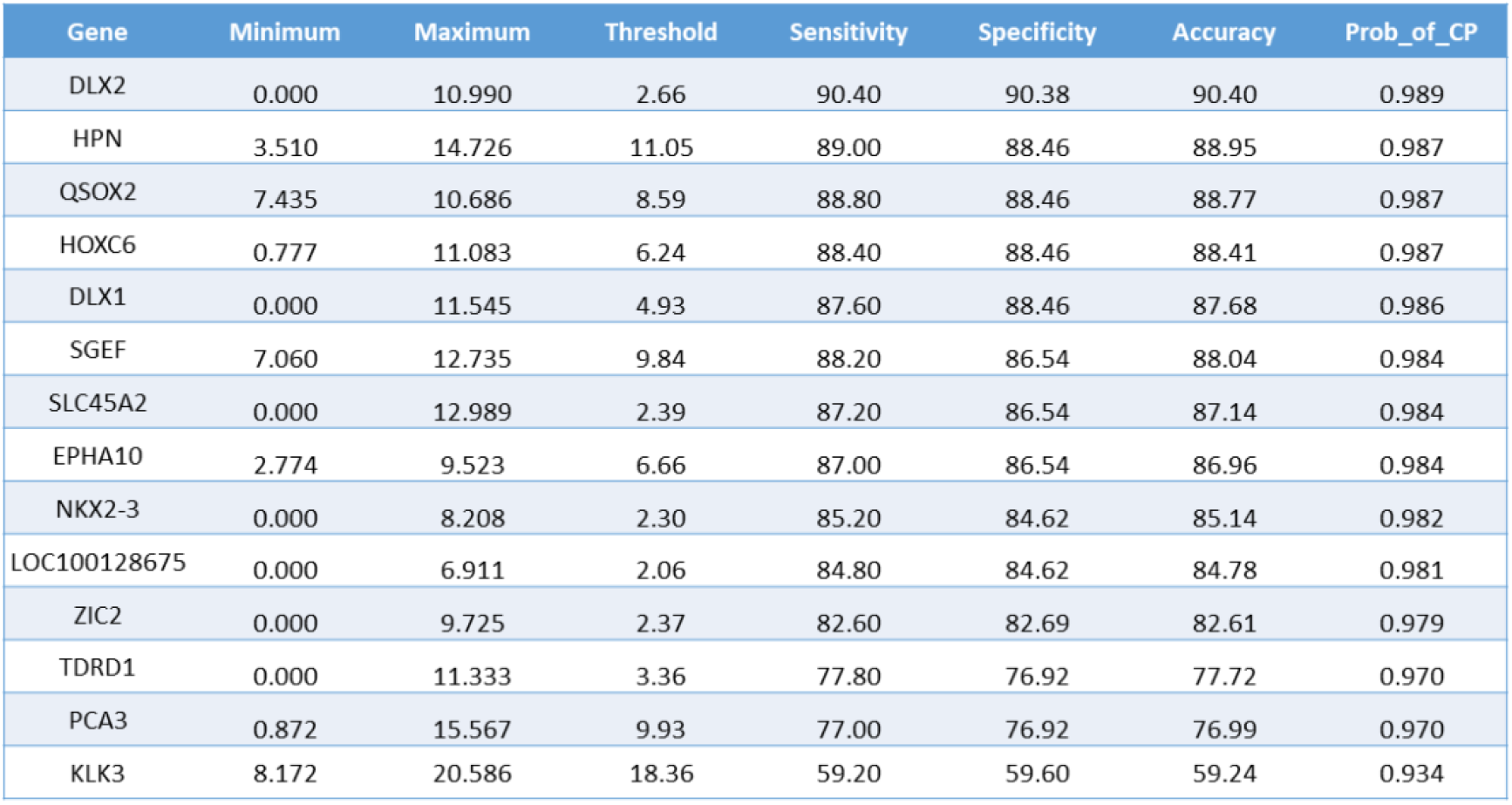
Ranking of genes based on their probability of correct prediction, the performance of each gene is computed using threshold-based approach.

In addition to ranking of genes, we also compute whether expression of these genes in prostate cancer samples is statistically significant or not. We plotted the gene expression value of the genes as box plot figures using GEPIA (Gene Expression Profiling Interactive Analysis) tool [17]. As shown in Figure 2, box plots generated depicts that 7 out of 14 genes (HPN, HOXC6, DLX1, SGEF, EPHA10, TDRD1, PCA3) are found to be significant.

## Discussion

In this study we aimed to identify gene expression-based biomarkers to distinguish between prostate cancer patients and a healthy control. We aimed to provide a tool to diagnose prostate cancer at an early stage. Patients with prostate cancer are usually diagnosed at a later stage of this disease as patients hardly develop any symptoms at an early stage of this cancer type. Better understanding of the molecular insights could aid in developing a tool which can detect the early onset of prostate carcinogenesis. We have extracted features using mean, significance difference in mean and AUROC based methods. We have identified the top ten genes based on these three methods and further applied machine learning techniques for classification of prostate cancer and healthy control subjects with high accuracy. In this study, we have converted expression values of top 10 genes identified by feature selection techniques to propensity index matrix. Classification models have been developed using propensity matric which showed significant improvement over the models developed using genes expression. This is a novel approach used in this study to improve the accuracy of correct prediction.

PSA being the widely accepted primary blood test for prostate cancer detection. We have also explored the possibility of KLK3 gene (PSA associated gene) as prostate cancer biomarker. Due to small difference between the mean expression value for normal and prostate cancer patient, KLK3 gene was not identified in the top10 genes in the feature selection techniques used in the current study. From figure 2 and 3, it is also evident that KLK3 gene performance as a biomarker for prostate cancer is not good in terms of sensitivity, specificity and accuracy. Various feature selection techniques were applied in order to determine the genes that can strongly distinguish between tumorous and non-tumorous records. It was observed that DLX1 and HOXC6 appeared to be in the top ten genes set in all three feature extraction techniques. In a recent study, researchers have reported a method SelectMDx which proposed HOXC6 and DLX1 as RNA based urine biomarkers in their study [11]. They have reported AUROC of 0.85 with 93% sensitivity, 47% specificity and 95% negative predictive value and the PCPTRC AUROC as 0.76 on the validation cohort.

**Figure 3:**
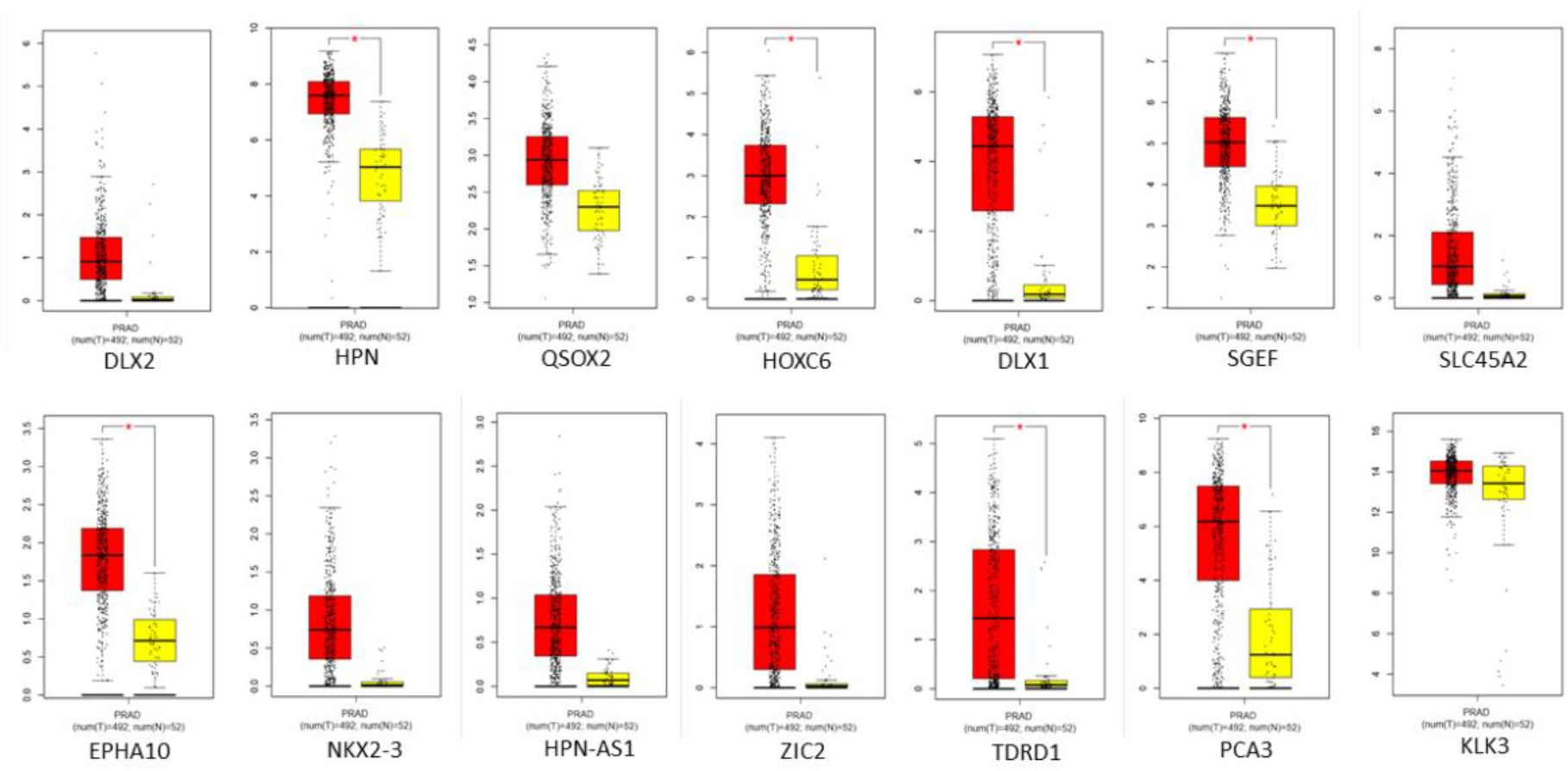
Box plots of ranked genes. In this figure red box plot depicts cancer samples and yellow depicts normal samples.

Another study reported a urinary three gene panel i.e. HOXC6, TDRD1, and DLX1 as a tool to distinguish prostate cancer patients with low sPSA values [10]. They have reported an AUROC value as 0.77 for these three biomarkers. We have also developed a model using these three biomarkers, ran the process ten times by shuffling the data each time, and achieved the average AUROC for KNN model as 0.927 ± 0.009 (Mean ± SD) for training and 0.971±0.002 for testing data set. We have also converted the expression values to propensity index and obtained the AUROC for SVC model as 0.981±0.002 for training and 0.914±0.001 for testing dataset. These results were highly unbalanced in terms of sensitivity and specificity.

In this study we have plotted Box plots to understand the potential of ranked top 14 genes based on probability of correct prediction. We have reported that HPN, HOXC6, DLX1, SGEF, EPHA10, TDRD1, PCA3 are found to be significant. Using past studies, we have mapped the role of DLX1, TDRD1 and HOXC6 in Prostate cancer. In literature, we have found studies which explains role of HPN [18], SGEF [19], EPHA10 [20] and PCA3 [8] in prostate cancer diagnosis and prognosis. These studies further validate our findings. In current study, we have applied 3 feature selection techniques i.e., mean based, standard deviation based and AUROC ROC based approach for identifying top 10 gene identifier for classifying prostate cancer sample vs. normal sample. Out of top 10 genes identified DLX1 and HOXC6 were found to be present using all three approaches. Further to understand the role of DLX1 and HOXC6 gene in functional pathways, we have explored GO annotations of both genes. DLX1 gene also known as Distal-Less Homeobox 1 is a protein encoding gene and is located on the long arm of chromosome 2. Gene Ontology (GO) annotation of DLX1 gene include sequence-specific DNA binding and chromatin binding. In literature DLX1 is reported to be associated with Dental Fluorosis and Witkop Syndrome. It is involved in related pathways such as DNA Damage/Telomere Stress Induced Senescence and Regulation of nuclear SMAD2/3 signaling pathway.

HOXC6 gene also referred as Homeobox C6 is also a protein coding gene and is located in a cluster on chromosome 12. Homeobox gene family usually encode a highly conserved family of transcription factors that are involved in a crucial role such as morphogenesis in multicellular organisms. Further Gene Ontology (GO) annotations include DNA-binding transcription factor activity and transcription corepressor activity. Diseases which are reported to be linked with HOXC6 include Lymphoma, Non-Hodgkin, Familial. With recent advancements, there is always a scope for improvement. Apart from these approaches, various other measures like entropy changes, etc. can be used to select the genes that will lead to higher information gain. We could also apply network analysis models to establish connections between various gene IDs. Building networks for tumorous and non-tumorous gene expression data could unfold deeper insights of the molecular mechanisms involved in development of the cancerous conditions.

## Abbreviations

PCa: Prostate cancer
GDC: Genomic Data Commons
TCGA: The Cancer Genome Atlas program
PRAD: *Prostate* Adenocarcinoma
PSA: Prostate Specific Antigen
sPSA: Serum PSA
PCA3: Prostate Cancer Antigen 3
DD3: Differential display code 3
AUROC: Area Under Receiver Operating Characteristics curve
MCC: Matthews Correlation Coefficient
Sens: Sensitivity
Spec: Specificity

## Ethics approval

‘Not applicable’

## Code availability

‘Not applicable’

## Author’s Contribution

### Conception and design

Shipra Jain, Kawal Preet Kaur Malhotra, and Gajendra P. S. Raghava

### Development of methodology

Shipra Jain, Kawal Preet Kaur Malhotra, and Gajendra P. S. Raghava

### Acquisition of data

Shipra Jain and Kawal Preet Kaur Malhotra

### Analysis and interpretation of data and results

Shipra Jain, Kawal Preet Kaur Malhotra, Sumeet Patiyal and Gajendra P. S. Raghava

### Writing, reviewing, and revision of the manuscript

Shipra Jain, Sumeet Patiyal and Gajendra P. S. Raghava

